# Congenital heart defects differ following left versus right avian cardiac neural crest ablation

**DOI:** 10.1101/2024.06.10.598133

**Authors:** Tatiana Solovieva, Marianne E. Bronner

## Abstract

The cardiac neural crest is critical for the normal development of the heart, as its surgical ablation in the chick recapitulates common human congenital heart defects such as ‘Common Arterial Trunk’ and ‘Double Outlet Right Ventricle’ (DORV). While left-right asymmetry is known to be important for heart development, little is known about potential asymmetric differences between right and left cardiac neural folds with respect to heart development. Here, through surgical ablation of either left or right cardiac neural crest, we find that right ablation results in more varied and more severe heart defects. Embryos with Common Arterial Trunk and with missing arteries occurred in right-ablated embryos but were not observed in left-ablated embryos; moreover, embryos with DORV and with misalignment of the arteries were more prevalent following right versus left cardiac crest ablation. In addition, survival of right-ablated embryos was lower than left-ablated embryos. Together, these data raise the intriguing possibility that there may be differences in left versus right cardiac neural crest during heart development.

## INTRODUCTION

Normal development of the heart requires an essential contribution from the cardiac neural crest. The cardiac neural crest is a migratory population of cells, first arising during neurulation, as the left and right neural folds meet in the dorsal midline of the embryo [1]. These cells then migrate through branchial arches 3, 4 and 6 [2] and through the truncus of the outflow tract toward the heart, laying down the truncal septum as they do so, thus dividing the outflow tract into two distinct vessels: the aorta and pulmonary trunk [3-5]. In addition to the outflow tract septum, cardiac neural crest has also been shown to contribute to the aortic arch arteries [3], subsequent carotid and other arteries [6] the ventricular septum [5], the cardiac ganglion cells [7] and some cardiomyocytes [8].

Importantly, ablation studies in the chick have demonstrated that absence of the cardiac neural crest gives rise to a range of heart defects [4, 9-13] reminiscent of common human congenital heart defects [14-15]. These include Common Arterial Trunk (also called ‘Persistent Truncus Arteriosus’, or PTA), ‘Double Outlet Right Ventricle’ (DORV), misalignment of arteries and abnormal myocardium function, among others. Having a four chambered heart, like mammals, the chick embryo is an appropriate model to better understand the specific roles of cardiac neural crest in congenital heart defects affecting humans [16].

While it is recognized that left-right asymmetry is important for heart development, the neural crest has generally been assumed to be a bilaterally symmetric population, given generally similar migration patterns on the left and right side of the embryo in cranial and trunk regions. Therefore, there has been little previous research on possible left-right differences in the neural crest. One notable exception was a study by Kirby and colleagues nearly 40 years ago, in which they ablated different unilateral portions of neural crest, including some sub-regions of the chick cardiac neural crest, and assessed subsequent effects on development [9]. They reported some variation in resulting embryonic defects, including some heart defects between equivalent left and right neural crest ablated embryos. A more recent study remarked briefly on the prevalence of different heart defects following left versus right cardiac neural crest ablation in chick [17] and another study in cavefish noted slight variations in gene expression between left and right cranial neural crest [18]. These data raise the intriguing possibility that left and right neural crest cells may have different roles in development at particular axial levels.

While the potential asymmetry between left and right cardiac neural crest populations is intriguing, the two previous studies on this topic report differing results following unilateral left/right neural crest ablations [9,17].

There are several key factors that likely contribute to these discrepancies. First, we now know that cardiac neural crest specification in chick occurs precisely at HH10 [19], while the studies in question have been done either on a much broader range of stages (HH8-10 [9]) or at slightly younger stages (HH9+ [17]), thus not accurately testing the specific effects of removing only mature fully specified cardiac neural crest. Second, the rostro-caudal extent of ablated neural crest tissue differs, with Kirby and colleagues ablating only a portion of the cardiac neural crest, missing the rostral portion [9], thus not testing the role of the entire left/right cardiac neural crest population. Third, the level of detail in the analysis was not always sufficient to determine all relevant morphological details associated with resulting phenotypes, and the extent of variation across embryos.

Here, we expand and build upon this to present a detailed, comprehensive report of the morphological effects of unilateral ablation of the entire left versus right pre-migratory cardiac neural crest on heart development in the chick.

## MATERIALS AND METHODS

### Embryos

Chicken embryos were obtained from Rhode Island Red hens (Petaluma Farm). Eggs were incubated horizontally in humidified incubation chambers (38 °C). Embryo staging was carried out according to Hamburger and Hamilton [20].

### Micro-surgery

All ablations were performed *in ovo*. Following horizontal incubation to HH10 (∼1.5 days), the embryo was first ‘lowered’ in the egg by removing 2-3 ml of albumen from the blunt end of the egg via a syringe. The egg was then ‘windowed’ by cutting a small (∼1-2 cm length) hole in the top of the shell. To provide contrast and help visualize the embryos, filtered black ink diluted in Ringer’s solution was injected directly below the embryo. Only embryos at HH10, specifically those with 9.5-10.75 somites (a range equivalent to just ∼1 hr 30 of developmental time), and displaying typical anatomy were used for ablations. A pulled glass needle was used to cut a small hole in the vitelline membrane directly above the region of interest. The needle was then gently used to make an incision along the lateral length of the cardiac neural crest on one side of the embryo, followed by incisions at the rostral, caudal, and medial sides of the unilateral cardiac neural crest. The rostral limit of removed neural crest was equivalent to the level of the mid-otic placode, while the caudal limit was equivalent to the caudal level of somite three. Detached cardiac neural crest tissue was then removed from the egg. Throughout surgery, the embryo was kept hydrated with Ringer’s solution. Post-surgery, a few drops of Ringer’s with diluted penicillin-streptomycin were added to prevent infection. The egg was then sealed with electrical tape and immediately incubated for a further 7 days, to HH32-35. After 7 days all eggs were opened. All embryos that had died at some point during the 7-day incubation were staged, if possible, before being discarded. HH22 was the earliest stage that could be reliably documented – embryos that had died before HH22 had disintegrated too much for reliable staging.

### Embryo processing

Surviving embryos were fixed in 4 % Paraformaldehyde in phosphate buffer overnight (4 °C), rinsed 3X in PBS and then washed overnight in PBS (4 °C). Embryos were equilibrated in 5 % sucrose in PBS overnight (4 °C), 15 % sucrose in PBS overnight (4 °C), and finally in 7.5 % gelatin/15 % sucrose overnight (38 °C). Embryos were then snap-frozen in molds using liquid nitrogen and stored at -80 °C for at least one overnight prior to cryosectioning. 16 μm serial sections were cut and collected onto Superfrost slides (Fisher, 1255015) (every section was collected). These slides were kept at room temperature overnight before subsequent processing/storage at -20 °C.

### Histology

To better visualize the morphological characteristics of sections, Hematoxylin and Eosin (H&E) staining was performed. To remove gelatin, slides were kept in 42 °C PBS for 15 min. Slides were then washed in PBS-triton (0.3 %) for 15 min at room temperature, followed by a 5 min PBS wash at room temperature on a rocker. Slides were dipped briefly in double distilled H_2_0 and dried on a hot plate. Slides were kept in Hematoxylin solution (Sigma MHS32) for 10 min, then washed for 10 min under running deionized water. Slides were kept for 1 min in Scott’s water followed by a brief dip in tap water, and then 3 min in Eosin solution (Sigma HT110216). Slides were then washed for 10 min under running deionized water and dried on a hot plate. Slides were mounted using DPX mounting medium (Fluka, 44581) and left overnight to dry.

### Imaging and image processing

Serial sections were imaged using a Zeiss imager MZ with a color camera. Images of serial sections were assembled for each embryo into one figure (Affinity Designer), facilitating the identification and analysis of 3-dimentional anatomical structures throughout the depth of the heart.

### Categorizing heart defects

Serial transverse cross-sections were analyzed systematically through the entire heart of each surviving embryo (n = 35 embryos).

#### Septation defects

were categorized based on location of the septal defect within the heart and alignment of the outflow tract relative to the ventricles. Location of septal defects was split into four categories: 1. Outflow tract septation defect distal to the semilunar valves; 2. Outflow tract septation defect at the level of the semilunar valves; 3. Septation defect at the root of the outflow tract and 4. Septation defect between the ventricles. At the level of the outflow tract roots, septation defects were further categorized according to the degree of communication between the left and right ventricles: (a) A complete communication between left and right ventricles or (b) A partial communication between left and right ventricles (partial/delayed fusion between the proximal outflow cushions). Defects in alignment of the outflow tract relative to the ventricles was assessed by systematically analyzing serial cross-sections and checking which ventricles the outflow tract vessels (aorta, pulmonary trunk or single outflow tract vessel), were overriding. For example, a phenotype exhibiting 1 and/or 2 would be categorized as ‘Common Arterial Trunk’ (PTA). A phenotype exhibiting 3 and/or 4 along with the outflow tract overriding the right ventricle, would be categorized as DORV, while presence of 3 and/or 4 without any misalignment of the outflow tract relative to the ventricles would be categorized as an interventricular communication.

#### Aortic arch artery defects

were categorized by analyzing the number of aortic arch artery-derived vessels contributing to the outflow tract and looking at the order of fusion of these vessels along the distal-to-proximal axis (their alignment to one another). By systematically going through serial transverse cross sections in a distal-to-proximal order, the aortic arch artery-derived vessels could be traced from the left or right sides of the embryo, thus revealing if left-or right-derived vessels were missing. The right pulmonary artery (RPA) which, together with the left pulmonary artery (LPA) gives rise to the pulmonary trunk, could be identified as missing in embryos where one right-derived artery was absent and where the pulmonary trunk was formed by a single, left-derived vessel (LPA).

## RESULTS

### Right ablated embryos have a lower survival rate than left ablated embryos

Pre-migratory left or right cardiac neural crest was ablated from embryos at HH10 (specifically embryos with 9.5-10.75 somites), which corresponds to a time-point immediately prior to the cells’ migration out from the neural folds (Fig. 1A). Following surgery, embryos were re-incubated for a further 7 days until they reached HH32-35, by which stage cardiac neural crest cells in a wild-type would have condensed to form the aorticopulmonary septum of the outflow tract [3-5], allowing septal defects such as DORV and Common Arterial Trunk to be detected.

**Figure 1:**
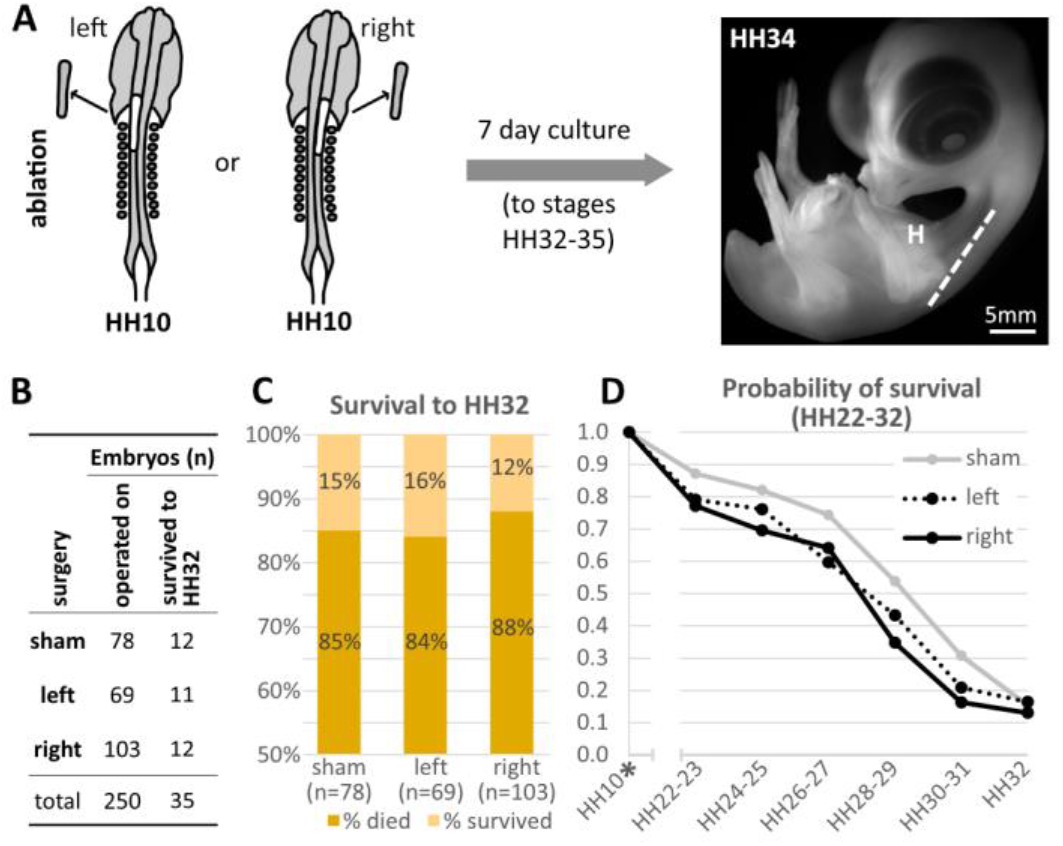
Experimental design and embryo survival. **A**. Experimental design for surgical ablations of unilateral cardiac neural crest. After 7 days of culture, embryos that survived ranged from stages HH32-35. Dashed line shows angle of sectioning. H = heart. **B**. Number of embryos operated on and number surviving each type of surgery/sham surgery. **C**. Survival of all embryos after 7 days of culture following surgery/sham surgery (to at least HH32). **D**. Survival probability (from HH22-32) of embryos undergoing left, right or sham ablations. Asterix at HH10 indicates the time at which surgery occurred.

Embryo survival to this stage was 16% for left ablated embryos (11/69 operated embryos), comparable to that of sham embryos (15%, 12/78 operated embryos) that did not undergo surgery but were prepared in an equivalent manner (window made in eggshell and hole made in the vitelline membrane above the embryo) (Fig. 1B-C). This suggests that most embryonic deaths that occurred were related not to the operation itself, but to the act of opening the eggs, and that at least by stage HH32, lacking a left cardiac crest is no more lethal to the embryo than having both cardiac neural crest populations intact. However, survival of right-ablated embryos was somewhat lower, at 12%, with only 12 out of 103 operated embryos reaching at least stage HH32, suggesting that the absence of right rather than left cardiac neural crest has more severe consequences on development during these stages. For embryos that died before HH32, stage at death was also recorded. This enabled the comparison of survival probabilities of embryos across the three different conditions (left, right, sham) (Fig. 1D). Probability of survival to all stages between HH22-31 was consistently lower for ablated embryos when compared to sham operated embryos, with right ablation resulting in slightly lower probability of survival than left ablation at almost all stages (Fig. 1D). The difference in survival probability reduces between the ablated and sham ablated embryos by HH32.

### Type and severity of septation defects varies between left and right cardiac neural crest ablated embryos

All operated embryos that survived to HH32-35 were embedded and sectioned (angle of whole-embryo sections shown in Fig. 1A) (Table 1, shows information on all surviving embryos). Serial cross-sections were analyzed through the entire heart and septal defects categorized at four levels: outflow tract vessels distal to the valves (Fig. 2A-Ei), outflow tract at the level of the valves (Fig. 2A-Eii), root of the outflow tract (Fig. 2A-Eiii) and through the ventricles (Fig. 2A-Eiv).

**Table 1:** All embryos. Information on embryo stage (at surgery and post-culture), observed cardiac defects and their locations for each processed embryo (n = 35).

| Embryo | Ablation type | Stage at ablation |  | Stage post culture | Septation defect | OFT septal defects |  | Interventricular communication |  | Arteries |  |
| --- | --- | --- | --- | --- | --- | --- | --- | --- | --- | --- | --- |
|  |  | Somite number | HH |  |  | connection of A and P | level | Type | level | number | fusion order |
| 310 | right | 9.5 | HH10 | HH34+ | Double outlet right ventricle | none | na | full | OFT roots | 5 | abnormal |
| 317 | right | 9.5 | HH10 | HH34 | partial | none | na | hairline | OFT roots | 5 | abnormal |
| 390 | right | 10 | HH10 | HH33+ | Double outlet right ventricle | none | na | full | OFT roots | 4 | abnormal |
| 411 | right | 10 | HH10 | HH33 | Common arterial trunk | yes | above OFT valves | full | OFT roots | 4 | abnormal |
| 413 | right | 10.25 | HH10 | HH34 | none | none | na | none | na | 4 | abnormal |
| 423 | right | 10.5 | HH10+ | HH33 | partial | none | na | hairline | OFT roots | 5 | normal |
| 435 | right | 10.25 | HH10 | HH34 | none | none | na | none | na | 5 | normal |
| 459 | right | 10.75 | HH10+ | HH32 | none | none | na | none | na | 5 | abnormal |
| 492 | right | 10 | HH10 | HH34+ | Double outlet right ventricle | none | na | full | OFT roots | 5 | normal |
| 504 | right | 10.5 | HH10+ | HH34+ | none | none | na | none | na | 5 | abnormal |
| 526 | right | 10.75 | HH10+ | HH34+ | Double outlet right ventricle | none | na | full | OFT roots | 5 | abnormal |
| 530 | right | 10.5 | HH10+ | HH33 | Common arterial trunk | yes | at level of OFT valves | full | OFT roots | 4 | abnormal |
| 242 | left | 10 | HH10 | HH32 | none | none | na | no | na | 5 | normal |
| 285 | left | 10.5 | HH10+ | HH32 | Double outlet right ventricle | none | na | full | OFT roots | 5 | abnormal |
| 299 | left | 10.5 | HH10+ | HH34+ | Double outlet right ventricle | none | na | full | OFT roots to ventricles | 5 | normal |
| 300 | left | 10.5 | HH10+ | HH34 | none | none | na | no | na | 5 | normal |
| 409 | left | 10.75 | HH10+ | HH33 | none | none | na | no | na | 5 | normal |
| 412 | left | 10.75 | HH10+ | HH34 | partial | none | na | hairline | OFT roots | 5 | normal |
| 414 | left | 10.5 | HH10+ | HH34 | partial | none | na | hairline | OFT roots | 5 | normal |
| 455 | left | 9.75 | HH10 | HH33 | partial | none | na | hairline | OFT roots | 5 | normal |
| 495 | left | 9.75 | HH10 | HH34+ | partial | none | na | hairline | OFT roots | 5 | normal |
| 503 | left | 9.5 | HH10 | HH32 | partial | none | na | hairline | OFT roots | 5 | normal |
| 516 | left | 9.5 | HH10 | HH33 | none | none | na | no | na | 5 | normal |
| Sham7 | sham | na | na | HH32+ | partial | none | na | hairline | OFT roots | 5 | normal |
| Sham18 | sham | na | na | HH32+ | none | none | na | none | na | 5 | normal |
| Sham27b | sham | na | na | HH34+ | none | none | na | none | na | 5 | normal |
| Sham32 | sham | na | na | HH34+ | none | none | na | none | na | 5 | normal |
| Sham37 | sham | na | na | HH35 | none | none | na | none | na | 5 | normal |
| Sham40 | sham | na | na | HH34 | none | none | na | none | na | 5 | normal |
| Sham44 | sham | na | na | HH34+ | none | none | na | none | na | 5 | normal |
| Sham51 | sham | na | na | HH33 | none | none | na | none | na | 5 | normal |
| Sham64 | sham | na | na | HH34 | none | none | na | none | na | 5 | normal |
| Sham68 | sham | na | na | HH33 | none | none | na | none | na | 5 | normal |
| Sham75 | sham | na | na | HH34 | none | none | na | none | na | 5 | normal |
| Sham78 | sham | na | na | HH33 | none | none | na | none | na | 5 | normal |

**Figure 2:**
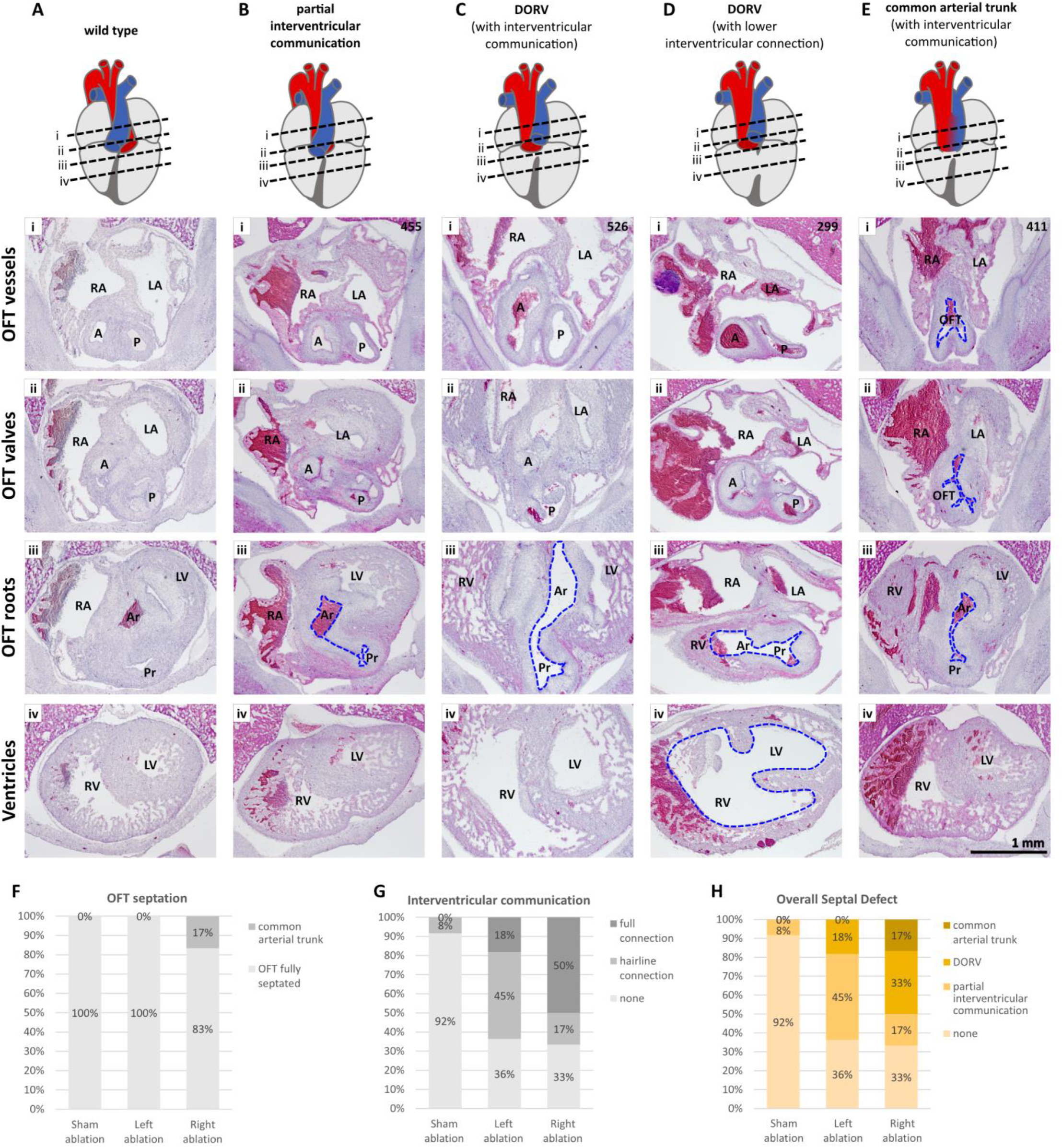
Septal defects. **A-E**. H&E serial cross-sections through hearts representing different heart defects. Corresponding diagrams of the hearts show levels for each section. Sections within a column come from a single heart. Sections from five hearts are shown. Blue dashed lines show communication between left and right sides of the heart that should not be present at this stage of development. **A**. Wild type. **B**. Partial interventricular communication. **C-D**. DORV. **E**. Common Arterial Trunk (PTA). **F**. Incidence of embryos with abnormal OFT septation. **G**. Incidence of embryos with full or partial interventricular communication. **H**. Frequency of defects affecting septation. A = aorta, P = pulmonary trunk, OFT = outflow tract, LA = left atrium, RA = right atrium, LV = left ventricle, RV = right ventricle, Pr = pulmonary root, Ar = Aortic root.

The complete absence of an aorticopulmonary septum through the outflow tract at, and distal to the level of the outflow tract valves is indicative of Common Arterial Trunk (also termed Persistent Trunk Arteriosus) (Fig. 2E). This cardiac defect is known to be the result of a failure of cardiac neural crest to successfully septate the outflow tract, resulting in a single outflow tract vessel rather than two separate vessels, the aorta and pulmonary trunk (Fig. 2Ei-ii versus Fig. 2Ai-ii). Intriguingly, Common Arterial Trunk was observed only following right cardiac neural crest ablations, occurring in 17% (n = 2/12) of right-ablated embryos (Fig. 2F). In these embryos, the single outflow tract was overriding the right ventricle.

The most common septal defect found was an interventricular communication (Fig. 2G), with a full connection between left and right ventricles at the level of the outflow tract roots (Fig. 2Ciii). Of these, in embryos with a separate aorta and pulmonary trunk, the aorta was misaligned, either partially or completely overriding the right ventricle rather than the left ventricle, suggesting DORV (Fig. 2C-D) [21]. DORV was observed in 33% (n = 4/12) of right ablated embryos and 18% (n = 2/11) of left-ablated embryos (Fig. 2H). In most observed cases of DORV, the interventricular communication was present only at the level of the outflow tract roots; however, in one case, there was a clear interventricular connection lower down in the ventricles of a left-ablated embryo (Fig. 2Div).

The third septal defect observed was that of a partial interventricular communication (Fig. 2B). This was characterized by the presence of a ‘seam’ or ‘hairline’ connection between the aortic and pulmonary roots, resulting from the incomplete (or delayed) fusion of the proximal outflow cushions, without complete septation *or* connection between the two sides (Fig. 2Biii compared to Fig. 2Aiii and Ciii). This phenotype was observed in 45% (n = 5/11) of left-ablated embryos and only 17% (n = 2/12) of right-ablated embryos (Fig. 2H).

Collectively, analysis of septal defects in the number of embryos available suggests more severe defects following right rather than left cardiac neural crest ablation.

### Artery development affected more severely by right than by left cardiac neural crest ablation

During development, cardiac neural crest contributes to the aortic arch arteries originating from branchial arches 3, 4 and 6, which subsequently undergo a significant remodeling. By HH32, at the levels studied, there are three arch artery-derived vessels originating from the right side of the embryo (aorta; right brachiocephalic artery, RBC; right pulmonary artery, RPA) and two from the left (left brachiocephalic artery, LBC; left pulmonary artery, LPA) (Fig. 3A). Serial cross-sections through these arteries were analyzed by following a series of distal-to-proximal ‘fusion’ events in the embryo (Fig. 3A-D).

**Figure 3:**
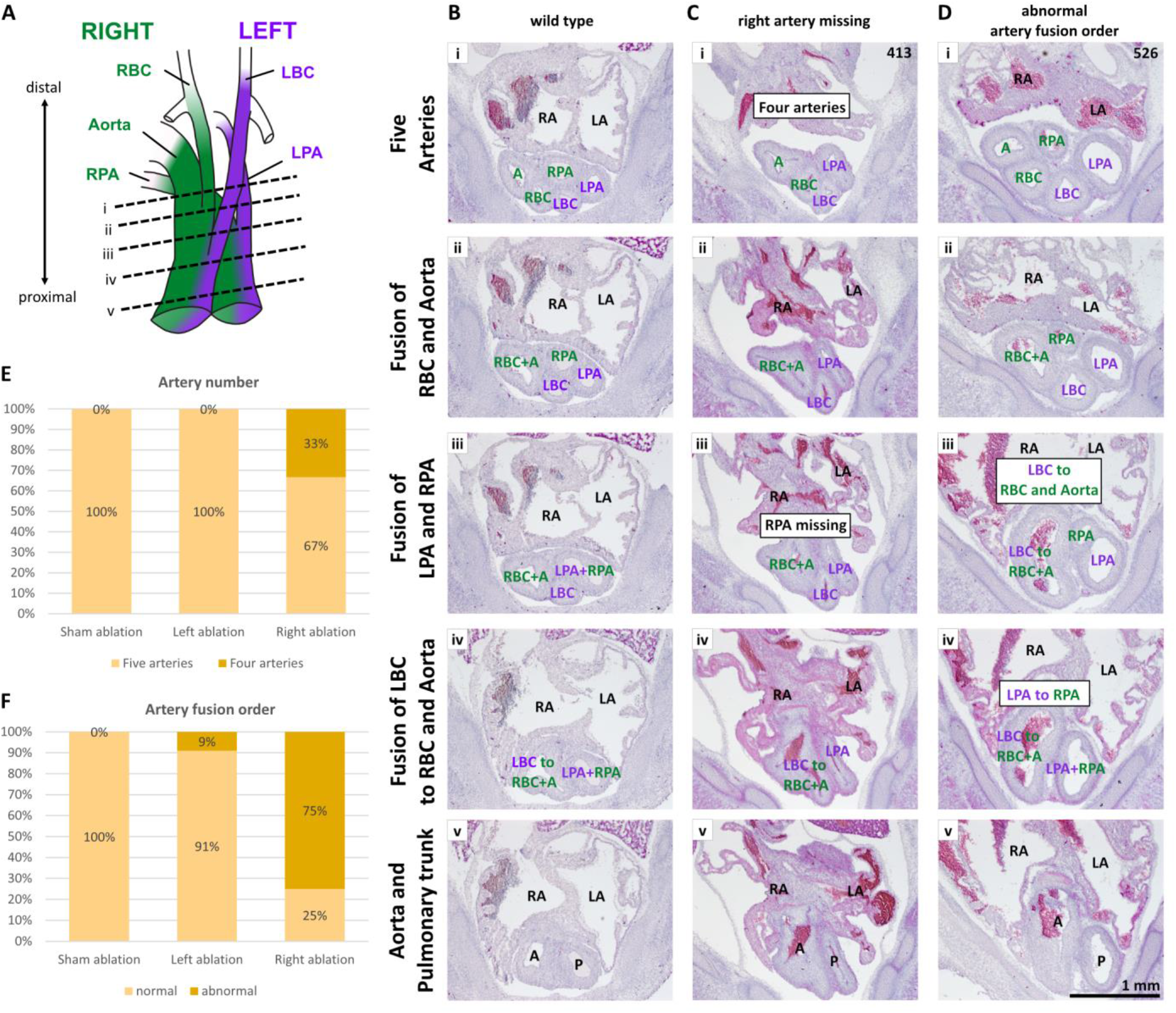
Artery defects. **A**. Diagram of arteries of the heart and their origins from the left/right side of the embryo, illustrating levels of sections shown in B-D. **B-D**. H&E serial cross-sections through the heart’s arteries, representing different defects in artery development. Sections within a column come from a single heart. Sections from three hearts are shown. **B**. Wild type. **C**. Artery missing. **D**. Abnormal fusion order of arteries along the distal-proximal axis. **E**. Incidence of embryos with a missing artery. **F**. Incidence of embryos with an abnormal artery fusion order. LBC/RBC = left and right brachiocephalic arteries, LPA and RPA = left and right pulmonary arteries, A = aorta, P = pulmonary trunk, LA = left atrium, RA = right atrium.

The most prominent defect observed was that of missing arteries (Fig. 3C). This occurred only following right cardiac neural crest ablation, in 33% (n = 4/12) of right-ablated embryos (Fig. 3E, Table 2). In these cases, there were only four arteries where there should normally be five (Fig. 3Ci versus 3Bi). In each case, a right-derived artery was missing, resulting in two instead of three arch artery-derived vessels originating from the right side of the embryo (right-ablated embryo with missing RPA shown in Fig. 3C).

**Table 2:** Embryos with missing vessels. Left/right branchial arch origin of missing vessels across embryos. LBC = left brachiocephalic artery, LPA = left pulmonary artery, RBC = right brachiocephalic artery, RPA = right pulmonary artery.

| Ablation | Embryos with missing vessels (n) |  |  |  |
| --- | --- | --- | --- | --- |
|  | Missing vessel<br>originating from<br>the left |  | Missing vessel<br>originating from<br>the right |  |
|  | LBC | LPA | RBC/Aorta | RPA |
| Right (n=12) | 0 | 0 | 1 | 3 |
| Left (n=11) | 0 | 0 | 0 | 0 |
| Sham (n=12) | 0 | 0 | 0 | 0 |

The second defect observed in the arteries following unilateral cardiac neural crest ablation was the order in which arteries fused to one another along the distal-to-proximal axis. In the wild type, it begins with the fusion of the RBC with the aorta, followed by the fusion of the LPA with the RPA (to form the pulmonary trunk) followed by the fusion of the LBC to the already fused aorta + RBC (Fig. 3A-B). The order, in which these fusion events occurred, but not the type of fusion events, was abnormal in 75% (n= 9/12) of right-ablated embryos and only in 9% (n = 1/11) of left-ablated embryos (Fig. 3F). For example, fusion of the LBC to the aorta and RBC occurred *before* fusion of the LPA to the RPA along the distal-proximal axis in the right-ablated embryo in Fig. 3D (Fig. 3Diii-iv versus 3Biii-iv).

Taken together, abnormal artery development appears to be almost an entirely right cardiac neural crest related phenomenon.

## DISCUSSION

Here, we have utilized unilateral cardiac neural crest ablations to study potential differences between the left and right cardiac neural crest during heart development and congenital heart defects therein.

The cardiac neural crest is a dynamic cell population that migrates from the neural folds through the branchial arches towards the outflow tract. The cells enter the aortic sac (HH25) and migrate towards the heart along the truncus of the outflow tract (HH26-31) laying down the aorticopulmonary septum as they do so, thus dividing the outflow tract into the aorta and pulmonary trunk. Complete septation of the outflow tract occurs by HH32 [5]. While survival probability of ablated embryos was consistently lower than that of sham embryos between HH22 and HH31, the difference in survival probability between these conditions declined by HH32. This timing (between HH31 and HH32) corresponds to a switch in the mechanism of outflow tract septation. Between HH26 and 31 the septation complex is made up of condensed, uncondensed and ‘scattered’ cardiac neural crest cells as they migrate proximally towards the heart. However, from HH31, the condensed portion of the septation complex ceases proximal migration. Instead, septation of the most proximal portion of the outflow tract occurs through the formation of layers of uncondensed cardiac neural crest cells [5]. This raises the possibility that reduced embryo survival may be linked with abnormal cardiac neural crest condensation. Furthermore, embryo survival between HH22 and 32 was almost always lower in embryos that underwent right versus left cardiac neural crest ablation. This would be in keeping with the finding that only right-ablated embryos displayed phenotypes with septation defects in the more distal levels of the outflow tract (e.g. Common Arterial Trunk, which is present at or above the level of the outflow tract valves).

In addition to cases of Common Arterial Trunk, right cardiac neural crest ablated embryos also had higher incidence of DORV and lower incidence of partial interventricular communication when compared with left-ablated embryos. Overall, our data suggest that right cardiac neural crest appears to cause more severe cardiac outflow tract septation defects than left cardiac neural crest. A caveat is that these observations are qualitative rather than quantitative given the small number of embryos and variation in phenotypes.

In relation to previous studies that performed variations of unilateral neural crest ablations, we are mindful of drawing direct comparisons because experimental design (stage at time of ablation, rostro-caudal extent of ablated tissue) and extent of analysis of resulting phenotypes can make direct comparisons difficult or, in some cases, inappropriate. Here we discuss some of the main similarities and differences across studies and the potential reasons behind any discrepancies.

The results in the present study for cases of Common Arterial Trunk (PTA) are comparable to those reported in Gandhi et al. [17], with a higher frequency following ablation of right cardiac neural crest. In contrast, in the study by Kirby and Colleagues [9], the frequency of Common Arterial Trunk was reported to be higher following left rather than right neural crest ablations (83% and 60% respectively) and overall, was much more common than in the present study. However, in the study by Kirby and Colleagues, a smaller portion of neural folds was removed (missing the rostral portion of the cardiac neural crest population) and the stages at which ablations were performed were much broader. Ablations were carried out between HH8-10 [9], a range which spans over 10 hrs of developmental time, compared to the present study where ablations were carried out at HH10, specifically on embryos with between 9.5-10.75 somites, which spans just ∼1 hr 30 of developmental time [20]. Between HH8-10, neural folds undergo significant changes [1], which may complicate interpretation of the resulting defects following ablation. The narrow developmental window used in the present study allows focus specifically on ‘mature’ cardiac neural crest after its specification is complete [19], and immediately prior to its migration away from, the neural tube. The study by Gandhi and Colleagues performed ablations at HH9+, a stage slightly younger than, but more comparable to the stages used in the present study, albeit analyzed in much less detail. Together these observations reinforce the case for right cardiac neural crest having a more significant role than left cardiac neural crest in the normal septation of the outflow tract.

In terms of ventricular septal defects, most embryos in the present study displayed some level of defect, with a comparable incidence in both right (67%) and left (63%) ablated embryos. However, following right ablations, the phenotypes were more severe (mostly full interventricular connections) as compared to those following left ablation (mostly partial interventricular connections). In reports by Kirby and Colleagues, ventricular septal defects were also present in most ablated embryos, but with a slightly higher incidence following left versus right neural crest ablations [9]. However, no details were available on the extent of ventricular septal defects (partial/full), thus complicating the comparison. In Gandhi et al., while incidence of ventricular septal defects was not explicitly reported, cases of DORV (which are always accompanied by an interventricular connection) were higher following left ablation [17]. This limited characterization of resulting phenotypes from both previous studies makes direct comparisons to the present study difficult. However, the overall tendency is that of a high incidence of interventricular communication following left *or* right cardiac neural crest ablation, with the present study suggesting a more severe phenotype following right rather than left cardiac neural crest ablation.

While previous studies of unilateral neural crest ablations have mentioned the presence of Common Arterial Trunk (PTA) [9, 17], interventricular septal defects [9] and DORV [17], to our knowledge, there are no previous detailed investigations into or findings of substantial differences in artery development following unilateral cardiac neural crest ablation. Here, a striking difference in development of arch artery derivatives was observed following the ablation of left versus right cardiac neural crest, with right ablation leading to artery loss and severely compromising correct artery alignment at stages HH32-35. In all cases, the missing artery was a vessel derived from the right side of the embryo, resulting in a phenotype with just two of the three right-derived vessels that are normally present. This raises the possibility that the prevalence of missing arteries following right versus left cardiac neural crest ablation might be linked to the extensive remodeling of aortic arch arteries and the asymmetric contribution of the subsequent vessel numbers from the left (two vessels) versus right (three vessels) sides of the embryo. One possibility is that removing the right cardiac neural crest simply resulted in insufficient cardiac crest cell number for all three right-derived vessels to develop. In contrast, the loss of left cardiac neural crest may not be detrimental to formation of only two left-derived vessels by stage HH32.

The finding from our study that removal of the right versus left cardiac neural crest has different effects on cardiac development raises the question of whether such developmental differences are the result of intrinsic differences in the left and right neural crest and/or a result of the different environments through which the cells migrate. For example, neural crest cells are known to interact closely with second heart field derived mesoderm/ myocardium as they migrate through the caudal pharyngeal arches and outflow tract [22, 23], thus raising the possibility that these cell types influence each other during development. It is possible that these influences may be different on the left and right sides of the embryo. For instance, the morphological and transcriptional asymmetries of the left versus right arch arteries [24, 25, 26] may affect left versus right cardiac neural crest differently as the neural crest cells migrate along these towards the heart. Further along their migration to the heart, behavior of left versus right cardiac neural crest might also be affected differently by the left/right asymmetries of the outflow tract, such as the ‘twist’ at the level of the conotruncal transition. It is therefore likely that the different effects of left versus right cardiac neural crest on heart development may result from a series of complex spatiotemporal interactions between cardiac neural crest cells, the second heart field and the asymmetrical environment though which they migrate, though intrinsic differences in the premigratory left versus right cardiac neural crest population may also play a role.

## CONCLUSIONS

In summary, in the present report we find that removal of the right pre-migratory cardiac neural crest results in more severe septation defects of the outflow tract and interventricular septum. Furthermore, right but not left cardiac neural crest ablation causes defects in artery number and alignment. Together, this raises the intriguing possibilities that left and right cardiac neural crest cells have intrinsic differences or that extrinsic factors differentially affect left versus right cardiac crest during heart development.

## ACKNOWLEDGEMENTS

We thank Prof. Robert H. Anderson for discussions on cardiac anatomy. This work was supported by grants from American Heart Association 1020751 to T.S., and NIH 1R01HL169287-01A1 and Additional Ventures 1011579 to M.E.B.

